# Molecular and Cellular Underpinnings of Spatial Heterogeneity in Fetal Cortical Folding

**DOI:** 10.64898/2026.01.20.700501

**Authors:** Xinyi Xu, Ruike Chen, Tianshu Zheng, Zhiyong Zhao, Mingyang Li, Dan Wu

## Abstract

Cortical folding is a defining feature of human brain development, yet the molecular and cellular mechanisms that produce regionally specific cortical folding remain incompletely understood. Here we combined high-resolution *in-utero* T2-weighted and diffusion MRI atlases (23-38 weeks of gestational age) with prenatal transcriptomic profiles to map regional macrostructural and microstructural features of cortical folding to underlying gene expression. We found genes whose expression patterns correlated with cortical curvature were enriched for neurogenesis, progenitor proliferation and radial glia guided neuronal migration, and localized to ventricular zone/subventricular zone progenitor cell subtypes, supporting for the protomap hypothesis of areal specification. By contrast, cortical microstructural markers-associated gene sets were implicated in myelination, cell adhesion and extracellular matrix remodeling, and mapped to astrocyte and endothelial cell programs. The microstructure-related gene expression peaked in the early postnatal period and remained high throughout childhood, while the curvature-associated gene expression reduced with age. Several of these cortical folding-related genes overlapped with autism spectrum disorder risk loci (e.g., *SCN2A, STXBP1*, *DVL3*). Collectively, these cross-modal findings outline a sequential developmental architecture—early progenitor driven patterning followed by myelin and extracellular matrix consolidation. In addition, we released a high-resolution fetal labeling atlas to facilitate further imaging-genetic studies of early cortical development.

## Introduction

Cortical folding is one of the most prominent and distinctive features of the human brain, characterized by the formation of gyri (ridges) and sulci (grooves) in the cerebral cortex. This intricate folding process is fundamental to the development of higher-order cognitive functions. Aberrant folding patterns have been associated with a variety of severe brain malformations and are implicated in numerous neurological, cognitive, and behavioral disorders, including autism spectrum disorder (ASD), epilepsy, and schizophrenia[1–5]. The most critical period for the emergence and rapid development of cortical folding (encompassing both morphology and microstructure) occurs during the fetal period, particularly between the second and third trimesters of gestation. Importantly, cortical folding exhibits pronounced spatial heterogeneity, with regional differences in curvature, sulcal depth, and the timing of gyrification across the cortical mantle[6, 7]. Previous postmortem brain studies have provided evidence that primary folds (such as sylvian fissure, central sulcus etc.) emerge first in a highly conserved spatiotemporal sequence across individuals, followed by secondary and tertiary folds[8, 9]. The consistent physical location and shape of primary folds suggest a genetic transcription influence on cortical folding[7, 10, 11].

During embryonic and early fetal development, intrinsic genetic mechanisms regulate key neurodevelopmental processes, including neurogenesis, neuronal migration, and neuroepithelial patterning[12–14]. These processes lead to the formation of transient neural structures such as the subventricular zone, preplate, and subplate, ultimately contributing to the establishment of the layers of the cortex and laying the foundation for cortical folding[12, 15, 16]. Various animal studies and ex-vivo human fetal studies have emerged to support hypotheses/models (e.g., protomap and protocortex hypotheses) explaining how distinct cortical areas form during development[14, 17–19]. However, we still lack a comprehensive in vivo overview of how complex molecular and cellular mechanisms of neurodevelopment give rise to the spatial heterogeneity of human cortical folding, primarily due to the challenges in in vivo imaging and molecular profiling of the human fetal brain.

Achieving a comprehensive understanding of human cortical folding demands methodologies that integrate molecular and structural analyses within intact tissue architectures. Recent advances in transcriptomic technologies now permit high-resolution mapping of gene expression across the human fetal cortex[20–22], allowing us to study early brain development at molecular level. Concurrently, improvements in *in-utero* fetal brain MRI, both in acquisition techniques and computational methods, have enabled non-invasive, high-resolution imaging of fetal brain morphology and microstructure[23–26]. Together, these tools offer unprecedented spatiotemporal precision for charting cortical folding processes from genotype to phenotype during the fetal window[27–31]. Consistent with this motivation, previous studies showed that regional variation in cortical gene expression aligns with structural and functional organization of the brain captured by MRI[32, 33], and that integrating spatially embedded adult human gene-expression atlases[34] with neuroimaging modalities bridges scale gaps[35] to reveal molecular correlates of structural and functional neuroanatomy, brain development, and disease[36–40].

In this study, we developed an integrative framework combining advanced *in-utero* fetal structural and diffusion MRI (dMRI) with transcriptomic profiling across mid-to-late gestation to elucidate the genetic, molecular, and cellular mechanisms driving cortical folding (Fig 1). Initially, we comprehensively characterized the regional variability of macro- and microstructures of fetal cortical folding, and identified genes whose expression patterns correlated this spatial heterogeneity. Next, we dissected the biological pathways, cell-type specificities and layer expression patterns of these candidate genes, and characterized their developmental expression trajectories, uncovering divergent molecular programs underlying macro-versus microstructural folding dynamics. We then assessed whether these cortical folding-related genes overlap with genetic risk factors for neurodevelopmental disorders such as ASD. Finally, to bridge molecular and anatomical perspectives and support imaging-genetic research, we released a high-resolution in-utero fetal brain atlas that aligned with the transcriptomic space for imaging-genetic analysis as an open-access resource.

**Fig 1.**
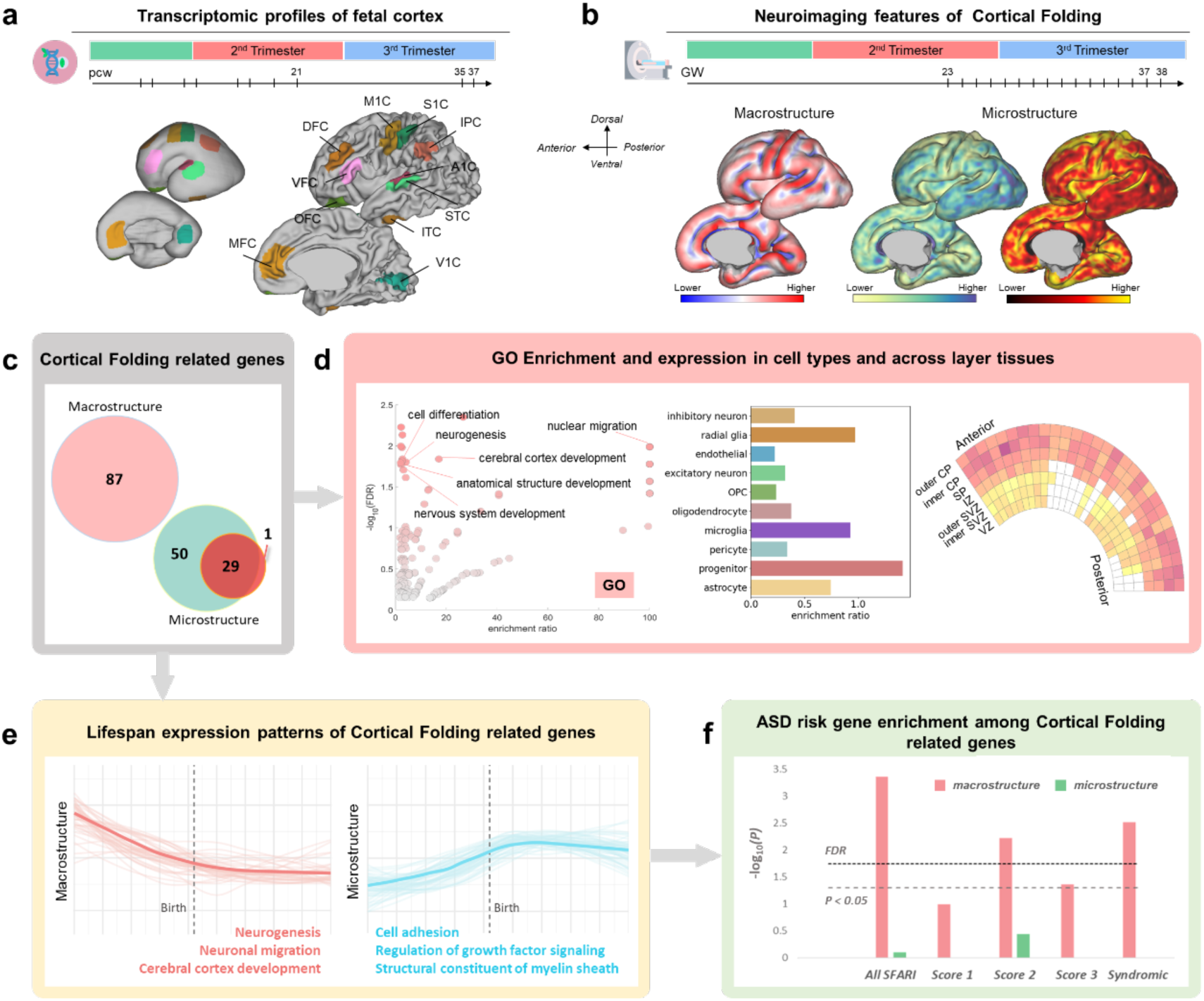
Overview of the current study. **(a) Transcriptomic resource.** Present study used prenatal bulk tissue mRNA data from PsychEncode project[22] to map spatial and temporal gene expression across the fetal cortex. (**b) Imaging phenotypes.** Representative cortical macro- and microstructural neuroimaging maps at 30 weeks of gestational age, which were used to characterize the regional cortical folding. **(c) Cortical folding related genes.** Genes associated with the spatial heterogeneity of cortical folding were identified through region-wise Pearson correlation analyses. (**d) Annotation.** Identified genes were characterized by enriched biological functions (left), cell type-specific expression (middle), and laminar expression patterns (right). (**e) Lifespan expression dynamics.** Nonlinear lifespan trajectories of cortical folding related genes, highlighting the cortical macrostructure-related genes that highly expressed prenatally (red) versus the microstructure-related genes that peaked postnatally (blue). (**f) Clinical relevance.** Overlap and enrichment of cortical folding related gene sets with curated autism spectrum disorder (ASD) risk genes from Simons Foundation Autism Research Initiative (SFARI) Gene database[41]. Abbreviations: pcw, postconceptional week; DFC, dorsolateral prefrontal cortex (PFC); VFC, ventrolateral PFC; MFC, medial PFC; OFC, orbital PFC; M1C, motor cortex; S1C, somatosensory cortex; IPC, posterior inferior parietal cortex; A1C, primary auditory cortex; STC, posterior superior temporal cortex; ITC, inferior temporal cortex; V1C, primary visual (occipital) cortex; GW, gestational week; GO, gene ontology; ASD, autism spectrum disorder; FDR, false discovery rate.

## Results

### Transcriptomic Signatures Underlying Spatial Heterogeneity of Cortical Folding

One central hypothesis underlying this study is that the temporal and regional dynamics of gene expression drive the observed morphological and microstructural heterogeneity of cortical folding. To test the hypothesis and understand the spatiotemporal relations, we first characterized both macro- and microstructural development of cortical folding across mid-to-late gestation using *in-utero* T2-weighted (T2w) MRI and dMRI. Leveraging our previously published 4D fetal brain T2w atlas[42] and dMRI atlas[43] (23-38 weeks of gestational age (GA) ; Supplementary Fig S1) in healthy fetuses, we computed curvature[44] and Diffusion Basis Spectrum Imaging (DBSI) model-derived microstructural parameters[45] (Fig 2a; Methods). DBSI is an advanced dMRI model that distinguishes microstructural elements, such as axons (fiber fraction), extracellular space (hindered fraction), cell body (restricted fraction), and free water[45–47]. These cortical folding features exhibited dynamic spatial variation and nonlinear developmental trajectories throughout the second and third trimesters (Fig 2a; Supplementary Fig S2-3; Supplementary Notes).

**Fig 2.**
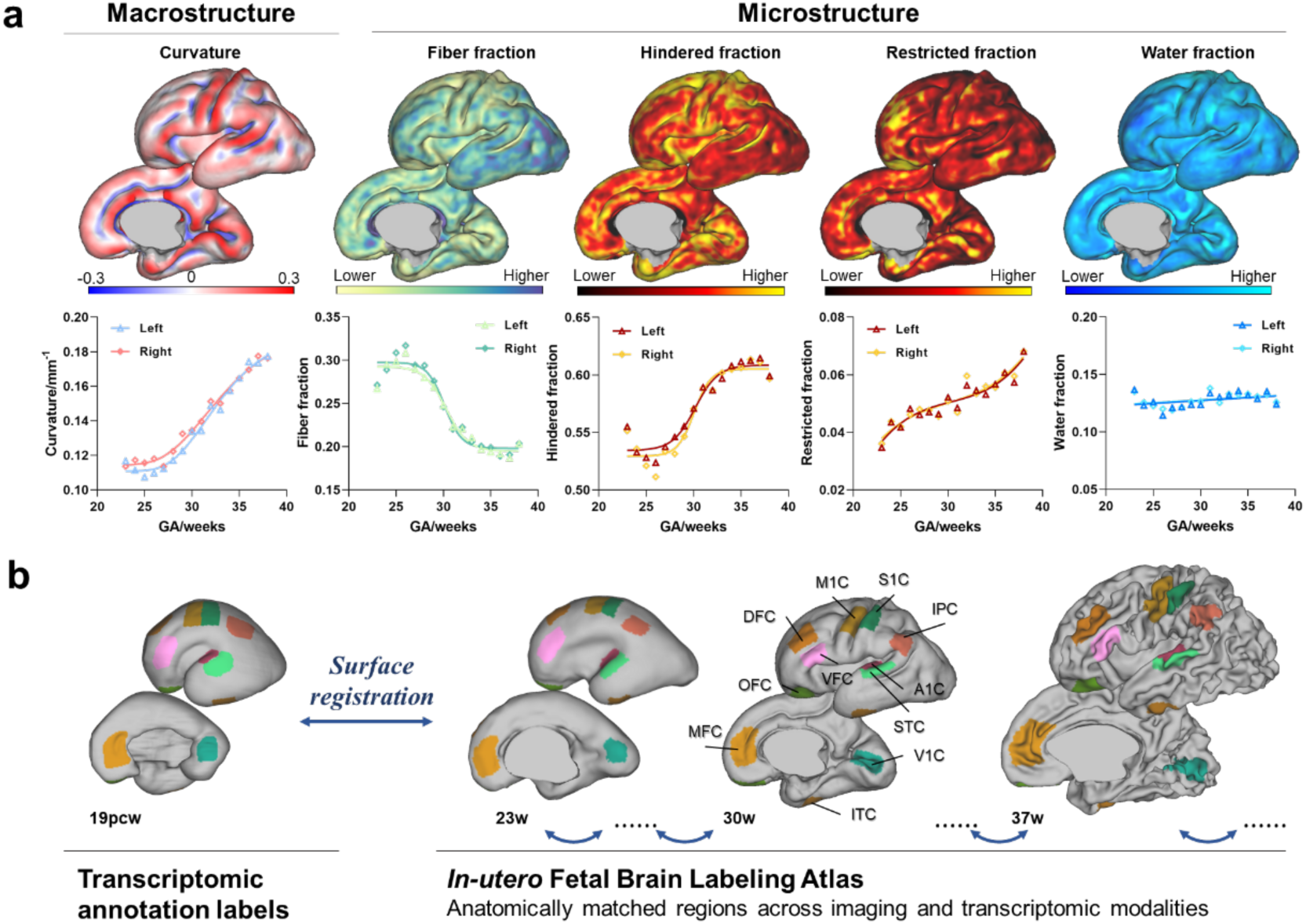
Spatiotemporal mapping of fetal cortical macro- and microstructural folding. **(a)** Top: Representative cortical surface maps of T2w-derived local curvature and diffusion-based DBSI metrics (fiber, hindered, restricted, free-water fractions) at 30 weeks GA, quantifying macroscopic and microstructural features of cortical folding. Bottom: Hemisphere-averaged trajectories for each imaging metric across the second-to-third trimester window (23-38 weeks GA). (**b)** Left: Developmental transcriptomic annotation labels from the PsychEncode project, mapping gene expression across 11 neocortical ROIs per hemisphere. Right: Fetal brain atlas with anatomically matched sample labels based on the fetal mRNA-seq atlas on the left, facilitating gene-imaging correlation analyses. Abbreviations: pcw, postconceptional week; w, week; DFC, dorsolateral prefrontal cortex (PFC); VFC, ventrolateral PFC; MFC, medial PFC; OFC, orbital PFC; M1C, motor cortex; S1C, somatosensory cortex; IPC, posterior inferior parietal cortex; A1C, primary auditory cortex; STC, posterior superior temporal cortex; ITC, inferior temporal cortex; V1C, primary visual (occipital) cortex.

Next, we utilized a developmental transcriptomic dataset of bulk tissue mRNA data sampled from 11 cortical regions in 4 prenatal brain hemispheres (the public PsychEncode project[22]; Table 1; Fig 2b) to link gene expression with cortical folding features. To enable precise alignment between imaging and transcriptomic data, we mapped PsychEncode anatomical labels onto our spatiotemporal atlas to generate the high-resolution fetal labeling atlas, thereby facilitating anatomically matched regions sampling across modalities (See Methods; Fig 2b). Following the methodology of Ball et al.[48], we selected 5,438 marker genes that were differentially expressed across cortical cell populations during gestation (Supplemental Table S1).

**Table 1.**
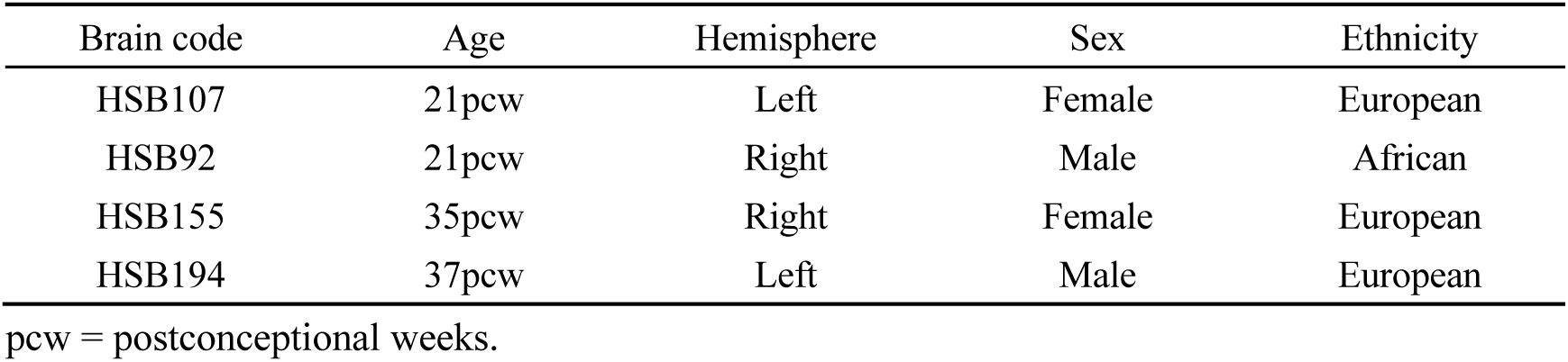
Included gene expression data from PsychEncode[22].

| Brain code | Age | Hemisphere | Sex | Ethnicity |
| --- | --- | --- | --- | --- |
| HSB107 | 21pcw | Left | Female | European |
| HSB92 | 21pcw | Right | Male | African |
| HSB155 | 35pcw | Right | Female | European |
| HSB194 | 37pcw | Left | Male | European |

pcw = postconceptional weeks.

We performed Pearson correlation analyses to assess the relationship between regional variation in gene expression and curvature across 22 anatomically matched regions of interest (ROIs) (i.e., 11 ROIs per hemisphere; Methods; Fig 3a). Out of the 5,438 genes, expression of 87 genes was spatially correlated with curvature at the early fetal stage (23 gestational week, corresponding to 21 postconceptional weeks transcriptomic data based on the terminology[49]) after correction for multiple comparisons (Supplemental Table S2). Of these, 45 genes exhibited positive correlations, indicating upregulated expression in regions with higher degree of cortical folding (denoted as *Curvature_up_*), and 42 genes exhibited negative correlations, i.e., down-regulated expression in regions with lower curvature (denoted as *Curvature_down_*) (Fig 3). Top correlated *Curvature_up_* genes included known molecular correlates of neural development (*IGFBPL1* - neurite outgrowth and axonal extension[50], *NEUROD4* - neural differentiation[51], *DVL3* - wnt signaling and neural patterning[52]*, SYNE2* - migration[53], *EOMES* - regional patterning[54], *DPYSL4* - axon guidance). Top correlated *Curvature_down_* genes were involved in neuronal migration and laminar organization (*YWHAH*), GABAergic neurotransmission (*GABBR2*), synaptic plasticity (*SYT5, PICALM*), and amyloid-β precursor protein modulation (*SPON1*). To validate these findings, we performed external validation using the publicly available fetal surface atlas developed by the Developing Human Connectome (dHCP) project[55] at 23w, and we identified 9 gene markers that were significantly associated with curvature, which were overlapped with our primary results (Supplementary Fig S4 and Table S2).

**Fig 3.**
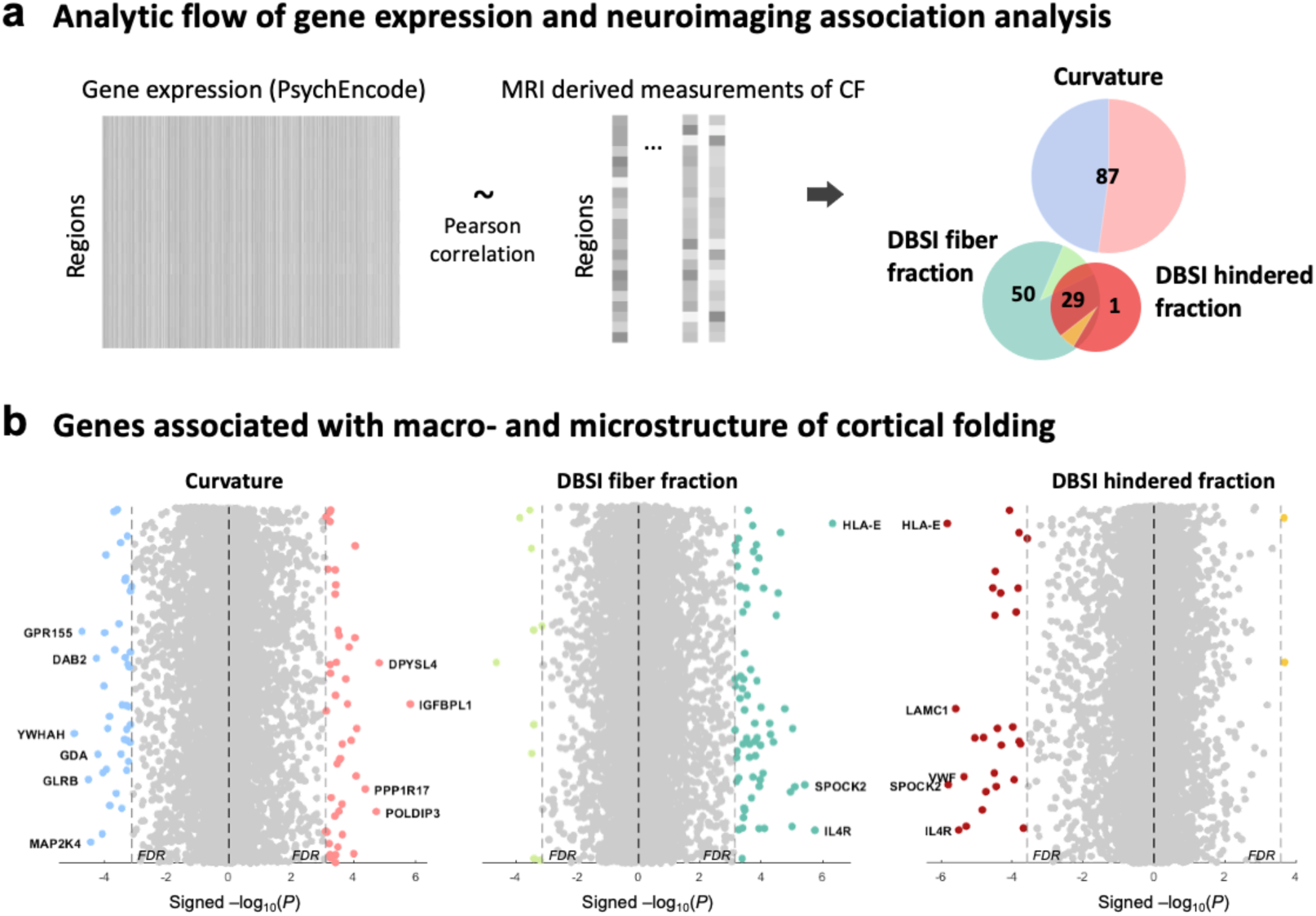
Analytic flow and summary results of the genes associated with spatial heterogeneity of cortical folding. **(a)** Associations between the spatial pattern of gene expression from the PyschEncode and MRI measurements of cortical folding were quantified using Pearson correlation analysis across 22 anatomically matched ROIs. (**b)** Results of Pearson correlation analysis of gene expression and cortical folding. Genes significantly associated with macro- and microstructure of cortical folding are marked with colors and top associated gene names are labeled.

Similarly, we identified 79 genes whose spatial expression patterns were significantly correlated with fiber fraction at late fetal developmental stage (37-38 gestational week), with 69 genes showing positive correlations (i.e., higher expression in regions with greater axonal/myelin density), hereafter denoted as *Fiber_up_* genes (Fig 3; Supplementary Table S3). Notable, *Fiber_up_* genes included oligodendrocyte differentiation and remyelination molecule (*GPR17*), neuroinflammation molecule (*PECAM1*) and cell adhesion molecules (*CD164)*. In addition, 30 genes exhibited significant associations with hindered fraction, 28 of which were negatively correlated, indicating elevated expression in regions of lower hindered fraction (denser cellular or fiber packing, reduced extracellular volume), and were termed as *Hindered*_down_ genes. Representative *Hindered_down_* genes included the well-known extracellular matrix organization molecule (*SPOCK2*), myelin-associated glycoprotein (*MAG*) and myelin basic protein (*MBP*) molecules - involved in axon regeneration and myelination[56], and cytoskeletal organization and cell adhesion regulator (*ACTN4*). Given the strong correlation between fiber and hindered fractions (*r* = −0.96, Supplementary Fig S5), 27 genes were shared between *Fiber_up_* and *Hindered_down_* gene lists (Fig 3a). Thereby, the following analysis focused on the *Fiber_up_* genes to avoid redundancy. The full gene lists, along with correlation coefficients and false discovery rate (FDR) values, are provided in Supplemental Table S3.

### Neurogenesis and neuronal migration underlie early fetal cortical folding

To elucidate the biological relevance of genes associated with curvature at mid gestation, we performed Gene Ontology (GO) enrichment analysis following FDR correction. Critical neurodevelopmental biological processes, including nuclear migration along microfilament (GO:0031022, enrichment ratio > 100, FDR = 0.0103), cerebral cortex radial glial guided migration pathways (GO:0021817, enrichment ratio > 100, FDR = 0.0167), neurogenesis (GO:0022008, enrichment ratio = 4.12, FDR = 0.0158), cell differentiation (GO:0030154, enrichment ratio = 2.74, FDR = 0.0145), were the top pathways enriched in *Curvature_up_* genes (Fig 4a top; Supplementary Table S4), indicating the fundamental roles of neuronal differentiation and radial glia-guided neuronal migration in shaping early cortical folding. In contrast, we found that genes highly expressed in less folded regions were significantly enriched in biological processes fundamental to synapse formation, function, and plasticity, including synaptic vesicle organization, neurotransmitter release, and calcium-dependent exocytosis (Fig 4a bottom; Supplementary Table S5). This result suggested that regions with relatively lower curvature may undergo active synaptogenesis and synaptic maturation.

**Fig 4.**
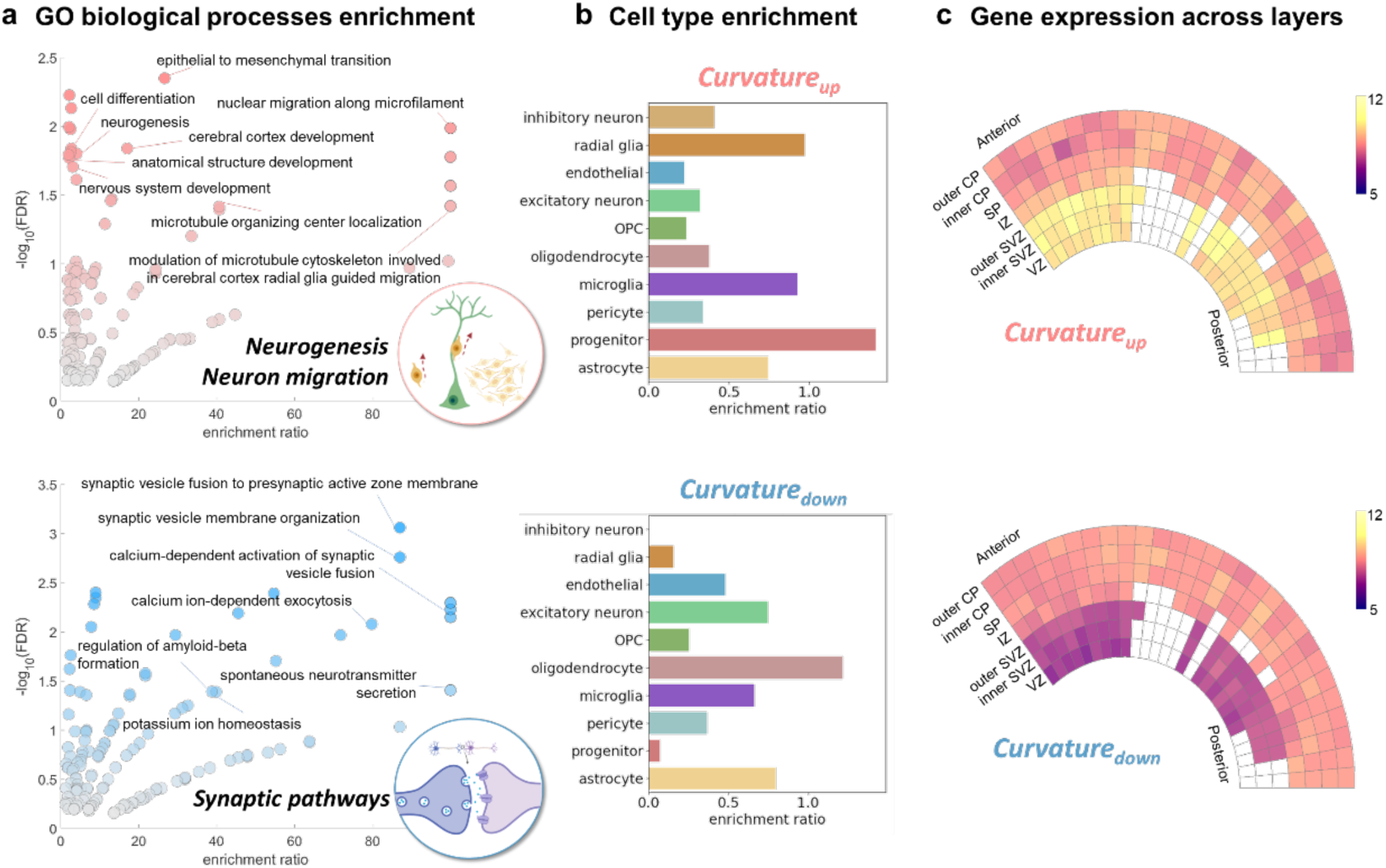
Macroscopic development of cortical folding is associated with neuronal and synaptic processes. **(a)** Volcano plots showing top representative biological processes of GO enrichment analyses in *Curvature_up_* (Top, pink) and *Curvature_down_* (Bottom, blue) gene sets identified at early fetal stage. (**b)** Fetal cell type enrichment for *Curvature_up_* (Top) and *Curvature_down_* (Bottom) genes. Bars show enrichment ratios for each fetal cortical cell class. **c.** Wedge plots show the pattern of gene expression across cortical layers. Rows correspond to cortical layers, and columns represent cortical regions ordered from anterior to posterior. Boxes are colored by gene expression level. Abbreviations: OPC, oligodendrocyte precursor cell; CP, cortical plate; SP, subplate; IZ, intermediate zone; SVZ, subventricular zone; VZ, ventricular zone.

We next found *Curvature_up_* genes were significantly overrepresented in progenitor populations (enrichment ratio = 1.4148, *P* = 0.0397; Fig 4b top; Supplementary Table S6; Methods). Within this cell class, post hoc analysis identified enrichment in several progenitor subtypes: excitatory neuron-like and radial glial-like progenitor cells[57] (both *P* < 0.01), medial ganglionic eminence-derived progenitors[57] (*P* = 0.012), and cycling progenitors in the G2/M phase[58] (*P* = 0.0489). Utilizing additional laser microdissection (LMD) microarray data from two fetal cortex samples at 21 postconceptional weeks (pcw)[21], we further observed higher *Curvature_up_* gene expression within the ventricular zone (VZ) and both outer and inner subventricular zones (SVZ) compared to postmitotic layers (cortical plate, subplate and intermediate zone; Fig 4c top). These findings are consistent with a growing body of evidence supporting the critical role of SVZ progenitor populations in driving cortical folding, i.e., continued production of upper layer neurons situated in the outer SVZ and expansion of outer SVZ radial glia lead to increased fold complexity in gyrencephalic species[59, 60], and manipulation of SVZ progenitor abundance modulates cortical folding in animal models[61–63]. By contrast, *Curvature_down_* genes expression was elevated within the postmitotic layers relative to germinal zones (VZ and SVZ; Fig 4c). This pattern further indicated that regions of lower curvature were characterized by a population of maturing, non-proliferative cells of glial lineages (e.g., oligodendrocytes and astrocytes).

Taken above findings together, we believe abundant neurogenesis and sustained neuronal migration of progenitors in the germinal zones, at least in part, support the early development of cortical folding during gestation, aligning with the established radial unit hypothesis whereby prolonged progenitor proliferation and radial migration yield a folded neocortex[64]. In this context, we reasoned that spatially differential expression of key transcriptional regulators likely modulates the intensity and duration of these processes across cortical regions, giving rise to the observed heterogeneity in folding patterns.

### Myelination, cell adhesion, and extracellular matrix remodeling accompany late cortical folding

Building on the early-stage patterns of exuberant neurogenesis and progenitor-driven folding described above, we next examined genes associated with microstructural metrics of cortical folding. Significant cortical microstructure-related gene associations were identified at the late fetal stage, a developmental window in which neurogenesis and migration have largely subsided, yet cortical folding continues toward an adult-like configuration. This period is characterized by rapid microstructural refinement[16].

Enrichment analysis of these late fetal cortical microstructure-related genes revealed significant overrepresentation of processes supporting microstructural development. *Fiber_up_* genes were enriched for GO biological process terms related to cell adhesion, fibroblast growth factor receptor signaling pathway, endothelial development, maintenance of blood-brain barrier, and apoptotic processes (Fig 5a; Supplementary Table S7). Moreover, the only significantly associated GO molecular function was structural constituent of myelin sheath (GO: 0019911, enrichment ratio = 75.85, FDR = 0.0366), with representative genes including *MBP, MAL, MOBP* (Fig 5b). Cell type-specific analysis indicated strong enrichment of astrocyte- and endothelial-expressed genes in the *Fiber_up_* genes (both *P* < 0.0001; Fig 5d; Supplementary Table S9). Together, these signatures implicate late-gestation myelination, cell adhesion, and axonal/fiber remodeling in establishing cortical microstructure and potentially in mechanically stabilizing emerging secondary and tertiary folds.

**Fig 5.**
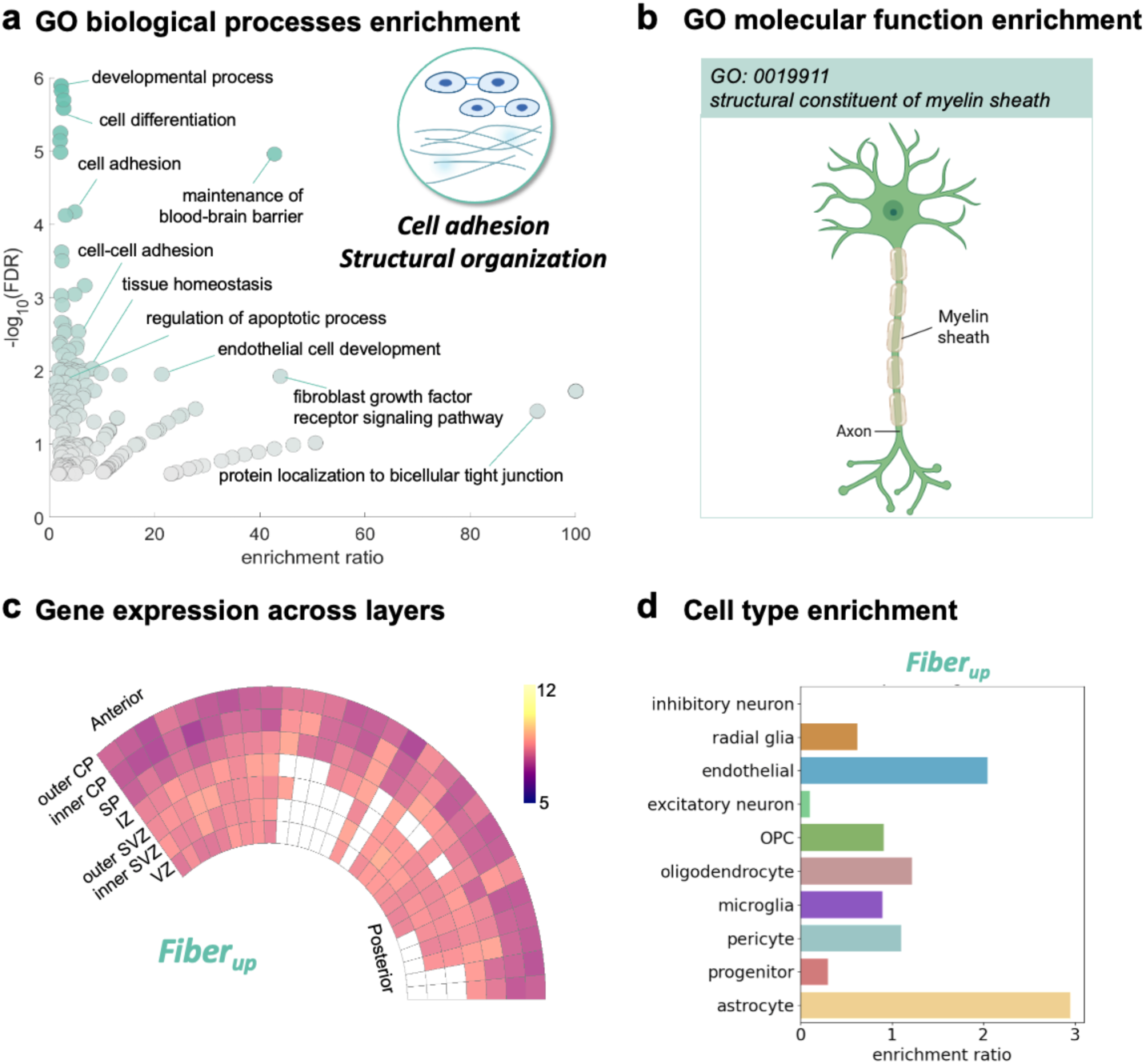
Microstructural programs associated with cortical folding (*Fiber_up_*). **(a)** Representative GO biological processes enriched among *Fiber_up_* genes identified at the late fetal stage. (**b)** GO molecular function enrichment for *Fiber_up_* genes. (**c)** Wedge plot for *Fiber_up_* genes shows the pattern of gene expression across cortical layers and regions. Rows correspond to cortical layers, and columns represent cortical regions ordered from anterior to posterior. Boxes are colored by gene expression level. (**d)** Enrichment ratio for fetal marker genes expressed by each cell class for *Fiber_up_* genes. Abbreviations: CP, cortical plate; SP, subplate; IZ, intermediate zone; SVZ, subventricular zone; VZ, ventricular zone; OPC, oligodendrocyte precursor cell.

In addition to cell-adhesion and blood-brain barrier maintenance, *Hindered_down_* genes were significantly enriched for extracellular matrix (ECM) organization and related processes (Supplementary Fig S6a; Supplementary Table S8). It is known that ECM remodeling may reduce extracellular void space and increase local tissue compactness, and may contribute to the microstructural differentiation of sulci versus gyri[65, 66]. Notably, using LMD microarray data from two 21pcw specimens, both *Fiber_up_* and *Hindered_down_* gene sets exhibited elevated expression level in subcortical layers (Fig 5c and Supplementary Fig S6a wedge plot), showing their potentially important roles in early white matter and subplate transient zones development during the fetal period.

### Distinct developmental profiles of cortical folding-related genes

The divergent and complex genetic architectures underlying cortical macrostructure versus microstructure phenotypes indicated these feature classes may follow distinct developmental programs. To better understand this, we further profiled the lifespan expression trajectories of cortical folding-related genes using postmortem data from PsychEncode (Fig 6a). Genes associated with cortical morphology (*Curvature_up_* and *Curvature_down_*) had high expression prenatally, followed by a decline after birth. In contrast, fiber-related genes reached peak expression around 4 months of postnatal age, which was consistent with the onset of brain myelination in healthy human development[67, 68], and then decreased slightly across later postnatal stages. These genes remained elevated relative to prenatal levels, consistent with continued maturation and refinement of microstructural programs after birth[16]. Given this temporal dissociation between the macrostructure and microstructure-linked genes, we further tested whether early macroscopic change predicts later microstructural maturation. Using structural equation modeling, we found that curvature changes at 23 weeks were significantly associated with fiber fractions measured at 35-38 weeks (Fig 6b, Table S10), indicating the initial brain shape may drive later microstructural differentiation[3, 69, 70].

**Fig 6.**
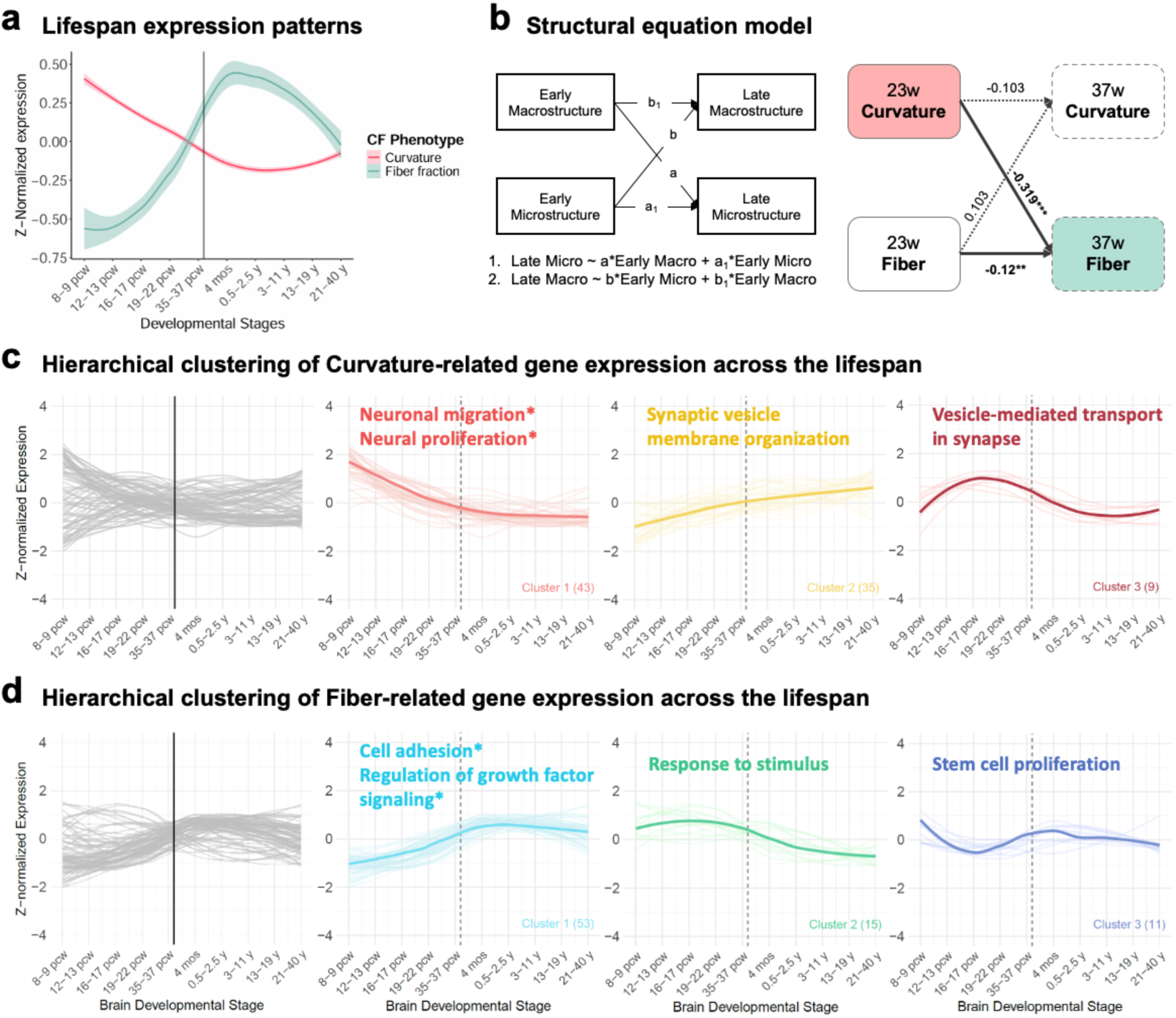
Developmental trajectories of cortical folding-related genes. **(a)** Lifespan expression trajectories (z-scored mean ± 95% CI) from PsychEncode bulk postmortem data for genes associated with macrostructural (curvature; pink) and microstructural folding (fiber fraction, green). (**b)** Structural equation model linking early macrostructural changes to later microstructural measures. Left: Path diagram. Right: A representative result of path coefficients and significance. Solid arrows indicate paths with FDR < 0.05; dashed arrows indicate paths with FDR ≥ 0.05. Significance notation: *, FDR < 0.05; **, FDR < 0.01; ***, FDR < 0.001. (**c)** Hierarchical clustering of curvature-associated genes (three clusters). Colored thick lines represent the average trajectory for each cluster, with the top representative biological pathways and the number of genes per cluster annotated. (**d)** Hierarchical clustering of fiber-associated genes (three clusters), with cluster means and top enriched pathways presented as in (c).

We next performed unsupervised hierarchical clustering on macro- and microstructural genes respectively (Supplementary Fig S7), and partitioned the genes into three principal developmental patterns (Supplementary Table S11). For the macrostructure set, the dominant cluster (cluster 1) was strongly enriched for pathways related to neuronal migration and progenitor proliferation, consistent with prominence of these processes prenatally (Fig 6c). In the microstructure set, the principal cluster (cluster 1) was enriched for cell-adhesion programs and processes involved in regulation of fibroblast growth factor signaling, which are key to maintaining fiber integrity and structure, reflecting postnatal microstructural consolidation (Fig 6d). Together, these findings indicated that the molecular substrates for microstructures and macrostructures of cortical folding follow distinct spatiotemporal programs with distinct neurobiological mechanisms.

### Cortical folding-related genes are associated with neurodevelopmental risk

Given their likely importance in shaping early brain organization, we hypothesized that cortical folding-related genes would be susceptible to severely disruptive mutations. Thus, we sought to investigate how these transcriptional programs may be affected in neurodevelopmental disorders. Here we overlapped the cortical folding-related gene sets (curvature-related genes and fiber fraction-related genes) with curated lists of rare ASD risk genes from the Simons Foundation Autism Research Initiative (SFARI) Gene database[41] (Fig 7a). Curvature-related genes showed significant enrichment for strong ASD candidates (SFARI score = 2) and for syndromic ASD genes (Fig 7b). Examples of overlapped curvature-specific ASD genes included the neuronal excitability regulator *SCN2A*, synaptic development factor *STXBP1*, radial glia scaffold integrity and migratory neurons regulator *LAMA1*, wnt signaling factor *DVL3* (See Supplementary Table S12 for the complete list of overlapping genes). Our findings provided evidence supporting that early disruptions in cortical folding may link molecular risk for ASD to the atypical cortical developmental patterns observed in affected individuals[71–73].

**Fig 7.**
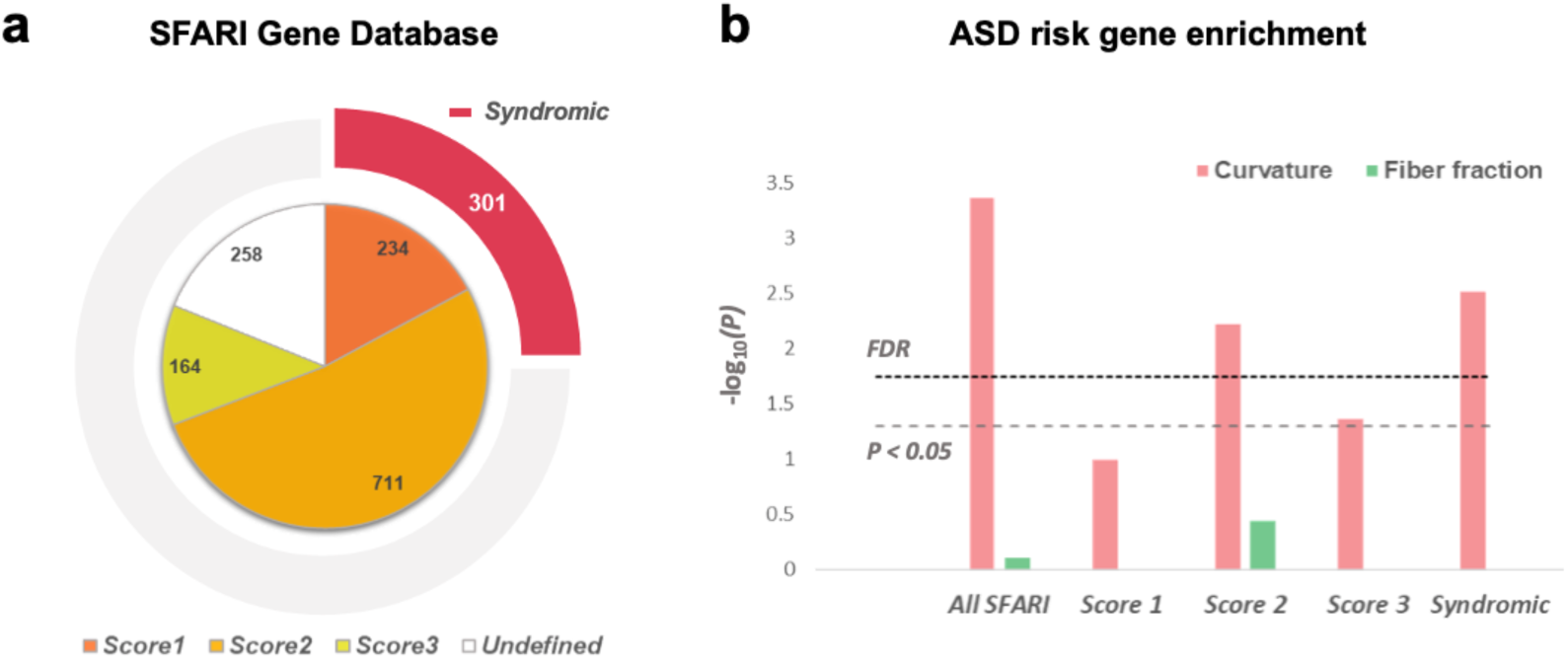
Enrichment of cortical folding gene sets in ASD risk genes. **(a)** Distribution of genes from the SFARI Gene Database, categorized by evidence strength: Score 1 (high confidence), Score 2 (strong candidate), Score 3 (suggestive evidence), and Syndromic (mutations associated with substantial risk and additional traits). (**b)** Enrichment of curvature-related (pink) and fiber fraction-related (green) gene sets within each ASD risk gene category. Significance is indicated by -log_10_(*P*) values, with FDR-corrected thresholds shown as dashed lines. Abbreviation: SFARI, Simons Foundation Autism Research Initiative.

## Discussion

In this study, we presented an integrative framework that combined *in-utero* MRI with prenatal transcriptomic data to non-invasively map the genetic, molecular, and cellular programs that determined the spatial heterogeneity of human cortical folding in vivo. Our multimodal analyses revealed two complementary developmental motifs. Early morphological (macrostructural) folding patterns were spatially coupled to progenitor-driven processes—proliferation, prolonged neurogenesis, and radial glia guided neuronal migration—consistent with radial unit type models of areal patterning and the establishment of initial cortical geometry. By contrast, microstructural differentiation of folds emerging later in gestation was associated with programs of myelination, cell adhesion and extracellular matrix remodeling, whose transcriptional signatures intensified around birth till early infancy. To facilitate further integration of molecular and imaging studies, we released a high-resolution fetal brain labeling atlas that maps the anatomical labels from transcriptomic data to the spatiotemporal cortical surface templates of our previously published in utero fetal brain atlases.

A central question in cortical development is how molecular and cellular programs shape the areal organization of the neocortex. Two models have been proposed to explain how distinct cortical areas are formed during development: the protomap hypothesis and the protocortex hypothesis[14, 64, 74, 75]. The protomap hypothesis posits that intrinsic molecular gradients in progenitor zones preconfigure areal identity and thereby bias subsequent cortical organization[64, 76]. By contrast, the protocortex hypothesis suggests that a uniform cortical sheet later acquires areal specialization through extrinsic input[75, 77]. Our results provide *in-vivo* human evidence consistent with intrinsic, progenitor driven areal patterning, by showing that regions of high curvature were enriched for progenitor- and migration-related transcriptional programs and that these programs localized to the germinal zones (VZ/SVZ) with regional variation. Experimental studies in gyrencephalic models and human tissue have shown that outer radial glia and intermediate progenitors are abundant in prospective gyral regions, and their disruption impairs folding[78, 79]. Moreover, waves of neurons generated by ventricular progenitors migrate along radial glia under the influence of transcription factors expressed in graded patterns, is proven to be a process essential for arealization[64, 80–83]. Our findings that *Curvature_up_* genes were enriched in progenitor populations, including outer radial glia and intermediate progenitors, and were preferentially expressed in the VZ and SVZ suggested regionally distinct proliferative dynamics early in gestation, which would help preconfigure the later morphological and functional architecture of the cortex. This molecular regionalization, evident before the establishment of sensory input or extensive synaptic activity, reinforces the view that intrinsic genetic programs drive early cortical patterning, consistent with the protomap framework[64, 84].

Another key finding is that cortical folding gene sets follow distinct, nonlinear developmental trajectories: macrostructure-linked genes peak prenatally and decline after birth, whereas microstructure-associated genes reach maximal expression in early infancy (∼4 months) before further postnatal modulation. This temporal dissociation implies sequential programs in which early molecular events (such as neurogenesis and neuronal migration) establish regional folding substrates, followed by later programs (such as cell adhesion and myelination) that consolidate and refine tissue microstructure. Consistent with this ordering, our structural equation modeling indicates that early curvature changes predict later development of microstructure. The results are also consistent with biomechanical models predicting that initial brain geometry biases subsequent gyrification[3], and with studies showing that cortical fold geometry influences white-matter microstructure[69, 70].

Atypical cortical development, particularly in local gyrification, has been repeatedly observed in individuals with ASD[71–73]. In our study, several cortical folding-associated genes overlapped with ASD risk genes, with the strongest enrichment seen for curvature-linked genes. For example, *SCN2A*, a *Curvature_down_* gene, encodes a voltage-gated sodium channel critical for action-potential initiation and neuronal excitability[85, 86]. Similarly, *STXBP1*, another *Curvature_down_* gene, regulates synaptic vesicle docking and neurotransmitter release[87, 88]. We also found ASD gene enrichment within the *Curvature_up_* set. For instance, *LAMA1*, a core laminin subunit of extracellular matrix and basement membranes[89], and *DVL3*, a wnt signaling mediator which controls progenitor proliferation, cell fate choices and tissue patterning[90]. While further experimental work is needed to establish causal links between cortical folding mechanisms and ASD pathology, our findings suggest that disruptions of curvature-related gene programs during critical fetal periods may interfere with cortical arealization and folding, offering potential molecular pathways through which ASD-associated cortical phenotypes could arise.

The combination of neuroimaging and transcriptomic technologies has created an unprecedented opportunity to identify the molecular, cellular, and pathway-level correlates of the imaging phenotypes. This combined approach has shown promise in revealing reliable underlying molecular, biological, and cellular pathways associated with brain structure and function[91, 92]. However, although promising, only a small number of studies have integrated fetal MRI with transcriptomic data to probe human prenatal brain development[93–95]. These pioneering studies delivered important insights but did not provide a complete portrait of cortical folding. For instance, Ball et al. focused on surface area expansion[93]; Lana et al. examined cortical-plate versus subplate thickness ratios[94]; and Hao et al. used ex vivo data to examine microstructural indices such as fractional anisotropy[95]. Moreover, these efforts lacked developmental continuity, limiting their ability to capture dynamic, stage-dependent changes over time. To bridge these gaps, the current study delivered a continuous, region-resolved mapping of both macro- and microstructural features of cortical folding to molecular and cellular signatures across second to third trimesters, and profiled the associated gene expression trajectories throughout lifespan.

However, several important limitations should be acknowledged. The molecular and imaging datasets were obtained from different populations, i.e., the transcriptomic resources are largely derived from European/African-ancestry donors, whereas the MRI atlases were constructed from a Chinese fetal cohort. We partially address this with a validation using the publicly available dHCP fetal surface atlas, but population-specific differences in gene expression or developmental timing may still influence some findings. Moreover, the PsychEncode fetal resource—while uniquely valuable—relies on relatively coarse spatial sampling, which limits our ability to resolve fine-grained spatial variation or to delineate precise boundaries between gyri and sulci areas. In addition, high-quality transcriptomic sampling across the full second-to-third trimester window remains sparse. We expect that forthcoming spatially resolved transcriptomic datasets, improved fetal brain imaging will enable more precise, longitudinal tests of the hypotheses generated here.

## Methods and Materials

### Fetal brain MRI data

#### Primary dataset – T2w atlas and dMRI atlas

We utilized our previously proposed fetal brain T2w atlas[42] and dMRI atlas data[43] spanning from 23-38 weeks GA, served as standard, unbiased average representations of a healthy fetal population (Supplementary Fig S1). In brief, these two atlases were generated from 90 and 89 healthy Chinese fetuses, respectively, using pair-wise registration initialization and iterative group-wise registration methods for each GA group. Further details on MRI acquisition protocols and image data processing can be found in methods sections of Xu et al.^[42]^ and Chen et al[43]. We chose to use population-average atlases for image-genetic analysis in this study to avoid the bias introduced by individual anatomical variability.

The T2w atlas data were processed using the dHCP-structural-pipeline (new-mirtk docker version, https://github.com/BioMedIA/dhcp-structural-pipeline) to reconstruct cortical surfaces, including white matter surfaces, midthickness surfaces, pial surfaces, sphere surfaces etc., and to extract the morphological characterization of cortical folding, specifically curvature[44]. Curvature was computed on white matter surfaces at each vertex to indicate the degree of cortical folding. Reconstructed surfaces and calculated curvature maps at each GA were visually inspected.

The dMRI data for each fetus were processed using the Diffusion Basis Spectrum Imaging (DBSI) model[45] to estimate the microstructural information of the fetal brain cortex. DBSI fitting was performed using the MATLAB application for DBSI calculations (https://osf.io/rmcjz/)[96]. The anisotropic fiber fraction was calculated and three isotropic component fractions were derived based on their difference in diffusivity (D), including restricted (0 < D ≤ 0.3 μm^2^/ms), hindered (0.3 < D ≤ 3 μm^2^/ms), and water (D > 3 μm^2^/ms) fractions. Other parameters were set as default. We derived parametric maps of fiber, restricted, hindered, and water components of brain tissue. These maps were then used to create the dMRI atlas of the four DBSI-derived parametric components, employing the same atlas generation method as described in [43]. Finally, these maps were further projected onto the cortical surfaces after co-registration with the corresponding T2w anatomical data at each GA.

#### Validation dataset – dHCP surface atlas

To further validate our findings, we used the publicly available spatiotemporal surface atlas generated by the Developing Human Connectome Project (dHCP) group, spanning from 21 to 36 weeks of gestation[55]. Specifically, we downloaded curvature maps for the 23-week GA data (the only time point that matches the age of transcriptomic data samples), and we aligned the white matter surface of the dHCP 23w GA atlas to our corresponding white matter surface atlas at 23 weeks using the curvature-based MSM_HOCR algorithm[97]. It is important to note that, unfortunately, the dHCP surface atlas does not include dMRI-based data, which limits our ability to compare the findings from microstructural information related to cortical folding.

### Bulk tissue transcriptome mRNA-sequencing data

In this study, we downloaded preprocessed bulk tissue cortical gene expression data obtained from the publicly available PsychEncode project (http://development.psychencode.org/)[22]. Briefly, tissue-level mRNA sequencing was performed on high-quality postmortem tissue samples from 16 anatomical brain regions (including 11 cortical areas of the neocortex: dorsal frontal cortex [DFC], ventral frontal cortex [VFC], orbitofrontal cortex [OFC], medial frontal cortex [MFC], primary motor cortex [M1C], primary somatosensory cortex [S1C], inferior parietal cortex [IPC], primary auditory cortex [A1C], superior temporal cortex [STC], inferior temporal cortex [ITC], and primary visual cortex [V1C], Fig 2b; and 5 subcortical areas: hippocampus, amygdala, striatum, mediodorsal nucleus of thalamus, and cerebellar cortex). These samples were collected from 41 specimens ranging in age from 8 pcw to 40 postnatal years[22]. Detailed anatomical boundaries for each cortical region at various developmental stages are provided elsewhere[20–22]. Gene expression levels were quantified as Reads Per Kilobase of transcript per Million mapped reads (RPKM).

Following the gene selection criteria outlined by Ball et al.[48], the initial dataset was filtered to include only protein-coding genes (NCBI GRCh38.p12, N = 18,453 out of 60,155 total genes from PsychEncode). To focus our analysis on genes expressed in the developing cortex, we further refined the list to include only those expressed in fetal cortical cells, based on a composite list of prenatal cell markers from five independent single-cell RNA studies on the developing fetal cortex[22, 57, 58, 98, 99]. This final selection comprised expression data for 5,438 genes (Supplementary Table S1). Gene expression values were log base 2-transformed after adding a pseudocount.

#### Cortical surface atlases with 11 marked ROIs

To facilitate comparison between prenatal mRNA gene expression data and fetal brain MRI data, we aligned the cortical ROI labels corresponding to the anatomical dissections used in mRNA analyses to our fetal brain MRI atlases. Anatomical annotations for the regional tissue dissections from PsychEncode[22], BrainSpan[21], Human Brain Transcriptome[20] projects were provided on a reconstructed cortical surface from a 19pcw specimen, as part of the Allen Institute BrainSpan Atlas of the Developing Human Brain. These cortical ROI labels were available for download (Fig 2b, https://www.brainspan.org/).

To propagate reference cortical ROIs to the MRI surfaces using automatic surface registrations, we further processed the downloaded data. First, we generated the corresponding spherical surfaces and calculated the curvature (both required for the subsequent surface registration) of the 19pcw cortical surfaces from BrainSpan Atlas using Freesurfer surface commands (https://surfer.nmr.mgh.harvard.edu/fswiki/FreeSurferWiki). Subsequently, the 19pcw cortical surfaces were co-registered to our atlas cortical surfaces at the earliest time point (23w GA) using multimodal surface matching with higher-order constrains (MSM_HOCR)[97]. Finally, The ROIs from the 19pcw surfaces were then transferred to our atlas cortical surfaces at 23w GA, based on the registration transformations, using workbench commands (https://www.humanconnectome.org/software/workbench-command).

For each gestational week, we applied this method to map 11 ROI labels per hemisphere onto the other cortical surfaces of our fetal MRI atlas. Visual inspection and manual refinement were performed to ensure consistency with the anatomical delineations described in[20–22]. This process resulted in a set of 11 cortical ROIs per hemisphere associated with regional bulk tissue mRNA data sampled during the prenatal period, co-registered with fetal neuroimaging data to enable correspondent sampling of cortical imaging metrics of fetal brain development. The ROIs maps, reconstructed surfaces, along with derived morphological and microstructural maps are available on https://github.com/zjuwulab/FetalFold-Atlas, facilitating imaging-genetics research on prenatal brain development.

### Correlation of gene expression and macro-/micro-structural measures of cortical folding

For this study, we selected RPKM data from 4 prenatal specimens (two at 21pcw, one at 35pcw, and one at 37pcw; Table 1) that matched best with the GAs of fetal brain MRI. Note that there is a two-week difference between pcw and GA according to the terminology[49]. We calculated the mean values of morphological (curvature) and microstructural (DBSI-derived fiber, hindered, restricted, and free-water fractions) measurements of cortical folding within each cortical ROI for both the left and right hemispheres. For each of the 5,438 selected genes, we constructed a vector representing the average expression levels across the 22 neocortex ROIs across the left and right hemispheres at the early fetal stage ([HSB107 – 21pcw Left, HSB92 – 21pcw Right]) and late stage ([HSB194 – 37pcw Left, HSB155 – 35pcw Right]), respectively. In parallel, MRI-derived measurements vectors were created using mean values from all cortical ROIs across the left and right hemispheres at early ([23w Left, 23w Right]) and late stages ([38w Left, 37w Right]). Note that, while the MRI measurements from the 39w fetal brain atlas were intended to match the 37pcw sample, the latest available GA in the atlas was 38w at the time of analysis.

We performed Pearson correlation analyses between each gene expression vector for the 5,438 genes and each cortical folding metric at the corresponding early and late fetal stages. To control for multiple comparisons, we applied the Benjamini-Hochberg false discovery rate (FDR) method.

### Enrichment analysis

To investigate functional and cellular enrichment of cortical folding-related genes, we performed the following analyses:

#### Gene Ontology (GO) enrichment

GO pathway enrichment was assessed using the PANTHER Overrepresentation Test available at https://geneontology.org/. Analyses were based on the GO Ontology database (DOI: 10.5281/zenodo.15066566, released 2025-03-16). Fisher’s Exact Test was applied with FDR correction for multiple testing. Unless otherwise specified, the Homo Sapiens genome was used as the reference background.

#### Cell-type enrichment

We tested whether cortical folding-related gene lists were over-represented among marker genes for major fetal cortical cell classes. Marker lists were derived from a previously published research[48]. Ten cell classes were evaluated: inhibitory neurons, excitatory neurons, radial glia, endothelial cells, oligodendrocytes, oligodendrocyte precursor cells (OPCs), microglia, pericytes, progenitor cells, and astrocytes. Enrichment was performed using an over-representation analysis (ORA) for each cell class, calculating the hypergeometric statistic as follows:

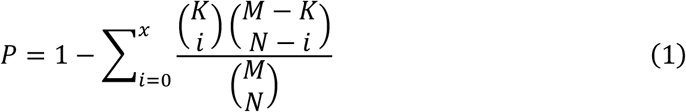

where *P* represents the probability of finding *x* or more genes from a target enrichment gene list - *K,* within a set of randomly selected genes - *N*, drawn from a background set - *M.* In this case, *N* refers to the gene list related to cortical folding, while *M* represents the total set of protein-coding genes. Enrichment ratios were calculated by comparing the proportion of genes linked to each cell type within our gene lists of interest to their proportion in the entire background set, defined as the full set of protein-coding genes:

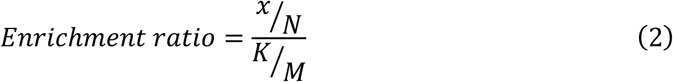

#### Enrichment for ASD risk factor

To explore how transcriptional gene programs related to cortical folding may be affected in neurodevelopmental disorders, we overlapped our identified gene lists with the Autism Spectrum Disorder (ASD) risk gene lists from the Simons Foundation Autism Research Initiative (SFARI) Gene database[41]. SFARI Gene is a comprehensive resource that ranks genes based on the strength of evidence linking them to autism risk. Genes are classified as follows: Score 1 (high confidence), Score 2 (strong candidate), Score 3 (suggestive evidence), and Syndromic (mutations associated with substantial risk and additional characteristics beyond the ASD diagnosis). We calculated hypergeometric *P* values for each overlap between our cortical folding-related gene sets and each SFARI score category using a similar ORA approach, by FDR correction for multiple comparisons.

### LMD Microarray data – layer specific expression pattern

To evaluate layer-specific expression of cortical-folding gene sets, we downloaded laser microdissection (LMD) microarray data from two 21-pcw donor brains, sampled across 29 cortical regions and 7 transient tissue layers from BrainSpan database (https://www.brainspan.org/). For details on tissue collection and dissection protocols, please see Miller et al[21]. For each identified gene set (i.e., *Curvature_up_*, *Curvature_down_, Fiber_up_,* and *Hindered_down_*), we computed the mean expression across genes and then averaged values across two donors. These mean, donor-averaged profiles were used to assess layer-enrichment and to generate the layer-resolved wedge plots shown in Fig 4c and Fig 5c.

### Developmental trajectories and hierarchical clustering

To characterize temporal expression patterns of cortical folding genes across prenatal and postnatal windows, we used the full PsychEncode lifespan dataset. Samples were binned into ten age windows: (1) 8–9 pcw, (2) 12–13 pcw, (3) 16–17 pcw, (4) 19–22 pcw, (5) 35–37 pcw, (6) 4 months, (7) 0.5–2.5 years, (8) 3–11 years, (9) 13–19 years, and (10) 21–40 years. Expression values were z-normalized across samples prior to analysis. For each imaging phenotype of cortical folding, the normalized expression values of all significant genes (FDR < 0.05) were averaged within each developmental window, and smoothed LOESS curves were plotted for visualization.

To identify dominant temporal expression patterns, genes were categorized into macrostructural (curvature-related genes) and microstructural (fiber fraction-related genes) groups, and unsupervised hierarchical clustering was applied. Pairwise distances between LOESS estimates were computed using Euclidean distance, and complete-linkage hierarchical clustering was applied to partition patterns into three clusters per group (cluster number chosen to balance interpretability and within-cluster coherence). Mean trajectories for each cluster were then computed for visualization. GO enrichment for each cluster was performed as described above; biological processes with adjusted *P* < 0.05 were considered significant.

### Structural equation modeling (SEM)

To examine directional influence between macrostructural cortical folding changes and microstructural alterations, we conducted structural equation modeling (SEM) in R (*lavaan* package). We examined these variables at early (23 weeks) and late (35-38 weeks) gestational stages. FDR correction was applied across four late gestational ages and four path coefficients.

## Data availability

The cortical labeling atlas, reconstructed surfaces, along with derived morphological and microstructural maps are available on https://github.com/zjuwulab/FetalFold-Atlas. The fetal brain T2w atlas and dMRI atlas data have been made publicly available on Github^1,2^. The dHCP surface atlas can be downloaded from https://doi.gin.g-node.org/10.12751/g-node.qj5hs7/. PsychEncode data are publicly available and can be downloaded at https://www.psychencode.org/data. LMD microarray data are available from BrainSpan project https://www.brainspan.org/. SFARI Gene database can be accessed at https://gene.sfari.org/tools/.

## Code availability

Code supporting data processing and analysis for this manuscript is available at https://github.com/zjuwulab/FetalFold-Atlas.

## Acknowledgement

This work was supported by the Ministry of Science and Technology of the People’s Republic of China (2021ZD0200202, DW), the National Natural Science Foundation of China (32427802 and U24A20754, DW), and Postdoctoral Fellowship Program of CPSF (GZC20251895, RKC).

## Supplementary Information

### Supplementary Notes

#### Regional Differences in Morphological and Microstructural Developmental Patterns of Cortical Folding

We first characterized the macro- and microstructural development of cortical folding across mid-to-late gestation using *in-utero* structural and diffusion MRI (dMRI). We leveraged our previously published fetal brain atlas data[1, 2], spanning from 23-38 weeks of gestational age (GA), as unbiased, healthy population-representative, pseudo-longitudinal templates (Supplementary Fig. S1).

As expected, both macro- and microstructural cortical folding metrics exhibited nonlinear developmental trajectories between 23 and 38 weeks GA (See trajectories in Fig. 2a; Supplementary Fig. S2). Total curvature increased gradually at first until around 27 weeks (Right hemisphere at 27.06 weeks; Left hemisphere at 27.69 weeks), then accelerated sharply, peaking in growth rate at around 31 weeks (Right - 31.49 weeks; Left - 31.47 weeks). Among microstructural measures, DBSI fiber fraction declined nonlinearly, hindered and restricted fractions increased nonlinearly, and free water fraction remained largely stable. Subtle hemispheric asymmetries were also apparent.

Regional surface maps of these phenotypes revealed dynamic spatial variations over gestation (Supplementary Fig. S2). To prepare for gene-imaging correlations, we then extracted 5 measures (curvature and DBSI fiber, hindered, restricted and water fractions) from 11 anatomically defined neocortical regions of interest (ROIs) per hemisphere, matched to the mRNA sequencing (mRNA-seq) data in a prenatal transcriptomic dataset (Fig. 2b; Supplementary Fig. S3)[3, 4]. In general, primary sulci and gyri exhibited higher curvature; gyral regions showed greater fiber fraction than sulci, whereas hindered fraction was elevated in sulcal cortex. Moreover, primary cortical areas (e.g., somatomotor cortex) displayed higher curvature and hindered fraction than higher-order regions (e.g., frontal, temporal lobes), while fiber and restricted fractions followed the opposite pattern (Supplementary Fig. S2 and S3).

### Supplementary Figures

**Fig. S1.**
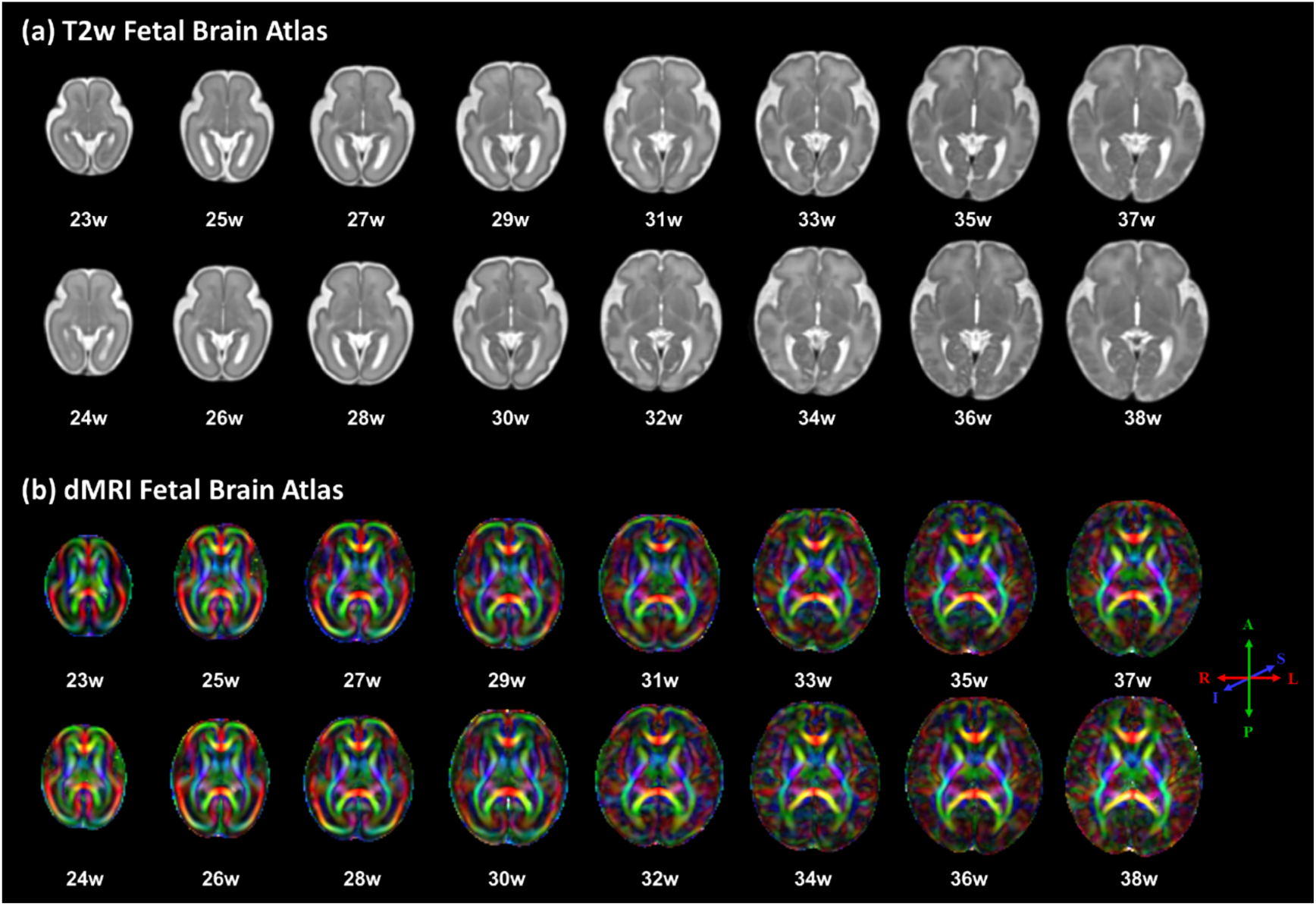
Spatiotemporal fetal MRI atlases (axial view). Axial slices from the structural (a) and diffusion (b) fetal brain MRI atlases spanning 23–38 weeks of gestational age, illustrating the templates used for surface reconstruction and Diffusion Basis Spectrum Imaging analysis to characterize cortical folding. Adapted from [1] and [2].

**Fig. S2.**
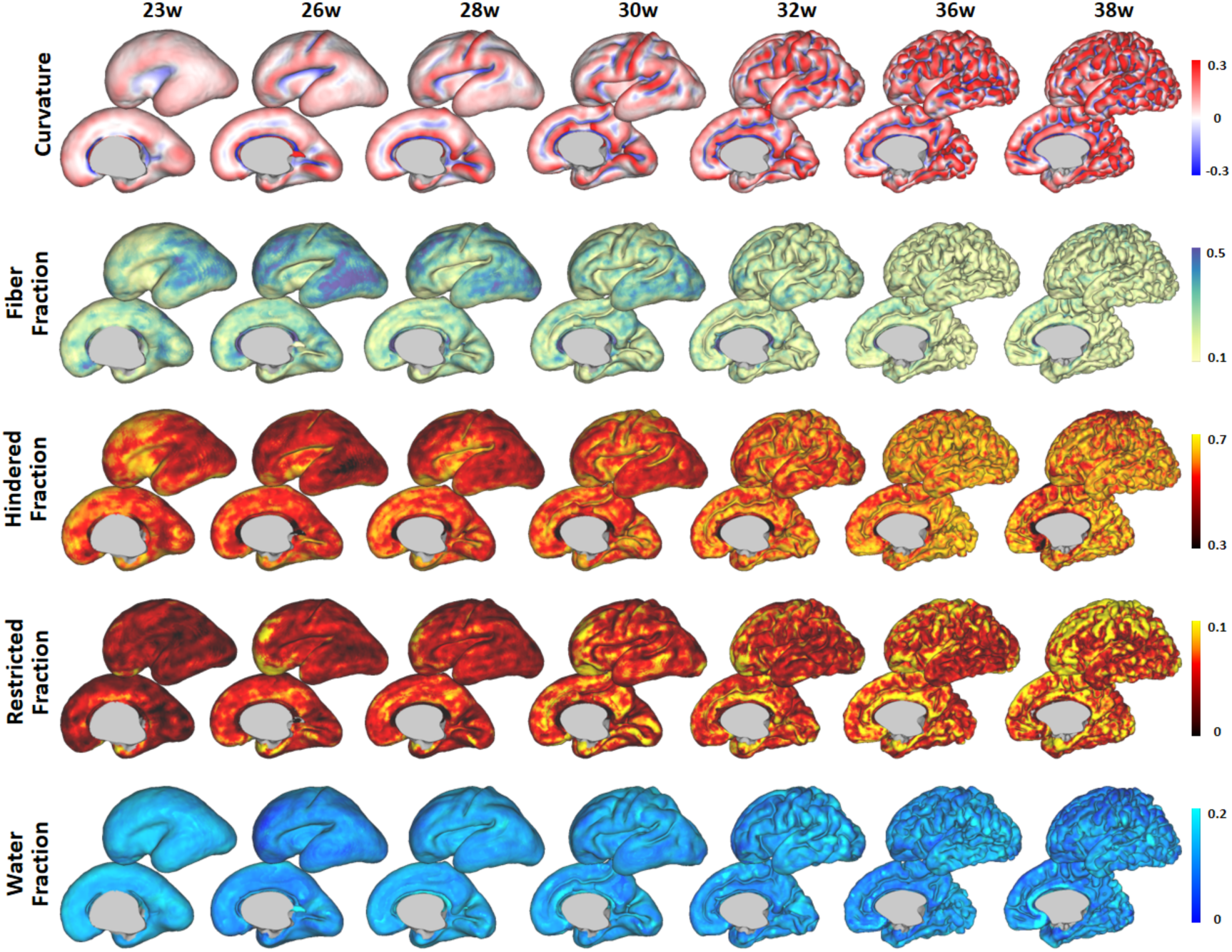
Weekly surface maps of macro- and microstructural folding (23–38 weeks of gestational age). Representative cortical surface maps of T2w-derived local curvature and diffusion-based DBSI metrics (fiber, hindered, restricted, free-water fractions) from 23 to 38 gestational weeks. These maps illustrate the spatio-temporal evolution of macroscopic folding and cortical microstructure.

**Fig. S3.**
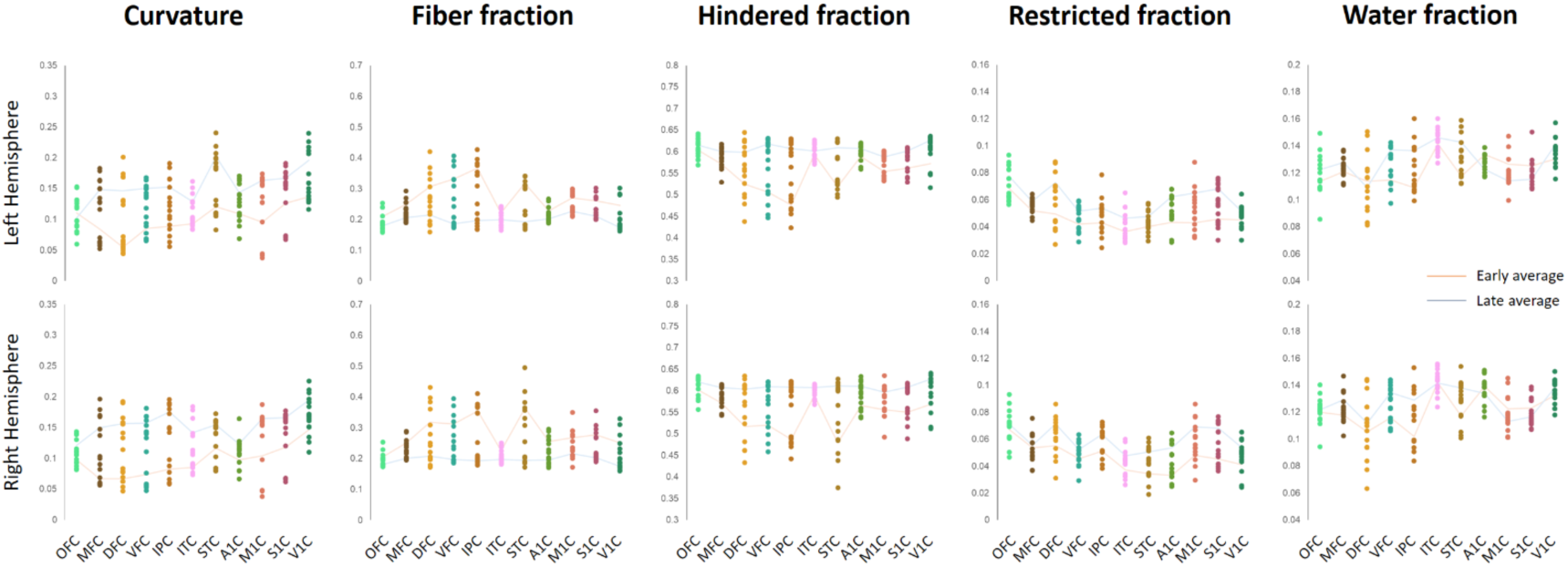
ROI-wise imaging phenotypes of cortical folding. Imaging phenotypes of cortical folding are shown for atlas at each gestational age within each cortical ROI (11 ROIs per hemisphere). From left to right: Curvature, DBSI fiber, hindered, restricted, and water fractions. Abbreviations: DFC, dorsolateral prefrontal cortex (PFC); VFC, ventrolateral PFC; MFC, medial PFC; OFC, orbital PFC; M1C, motor cortex; S1C, somatosensory cortex; IPC, posterior inferior parietal cortex; A1C, primary auditory cortex; STC, posterior superior temporal cortex; ITC, inferior temporal cortex; V1C, primary visual (occipital) cortex.

**Fig. S4.**
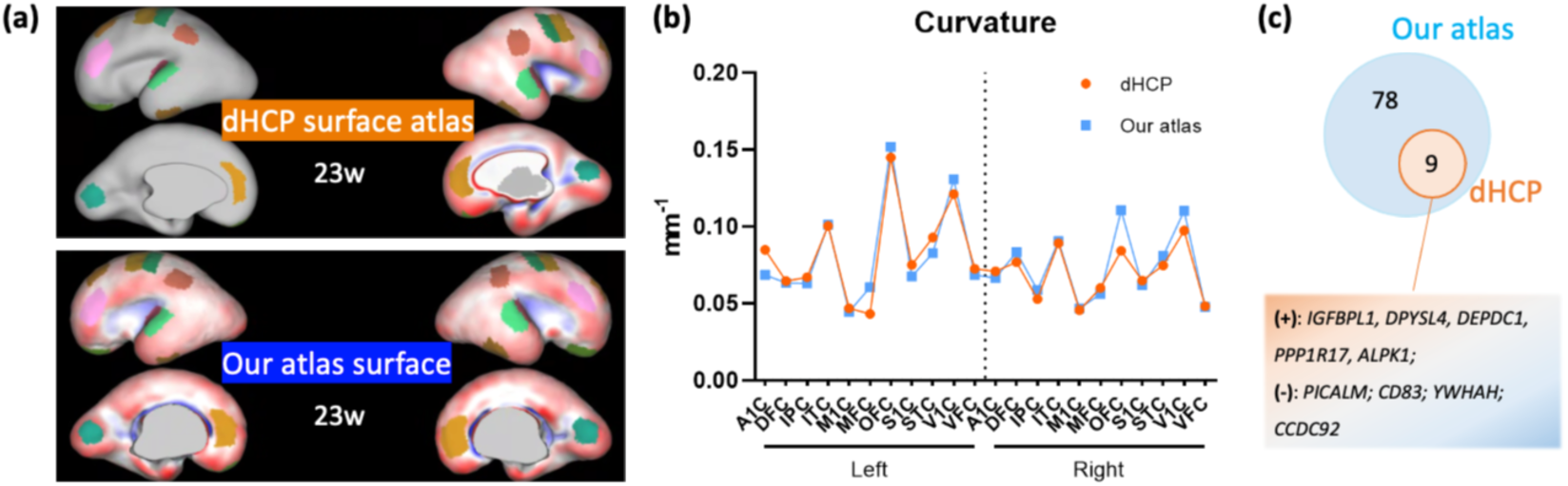
Validation using the dHCP fetal surface atlas. (a) Visualization of white-matter surfaces from the dHCP 23-week fetal atlas (top, orange) and from our atlas (bottom, blue). The red-blue overlay denotes local curvature, and the 11 cortical ROIs per hemisphere are indicated. (b) Line plots of mean curvature for the 11 ROIs in the left and right hemispheres (orange, dHCP; blue, our atlas). (c) Venn diagram summarizing gene-imaging association results; “+” indicates positive correlations and “−” indicates negative correlations. Abbreviations: DFC, dorsolateral prefrontal cortex (PFC); VFC, ventrolateral PFC; MFC, medial PFC; OFC, orbital PFC; M1C, motor cortex; S1C, somatosensory cortex; IPC, posterior inferior parietal cortex; A1C, primary auditory cortex; STC, posterior superior temporal cortex; ITC, inferior temporal cortex; V1C, primary visual (occipital) cortex

**Fig. S5.**
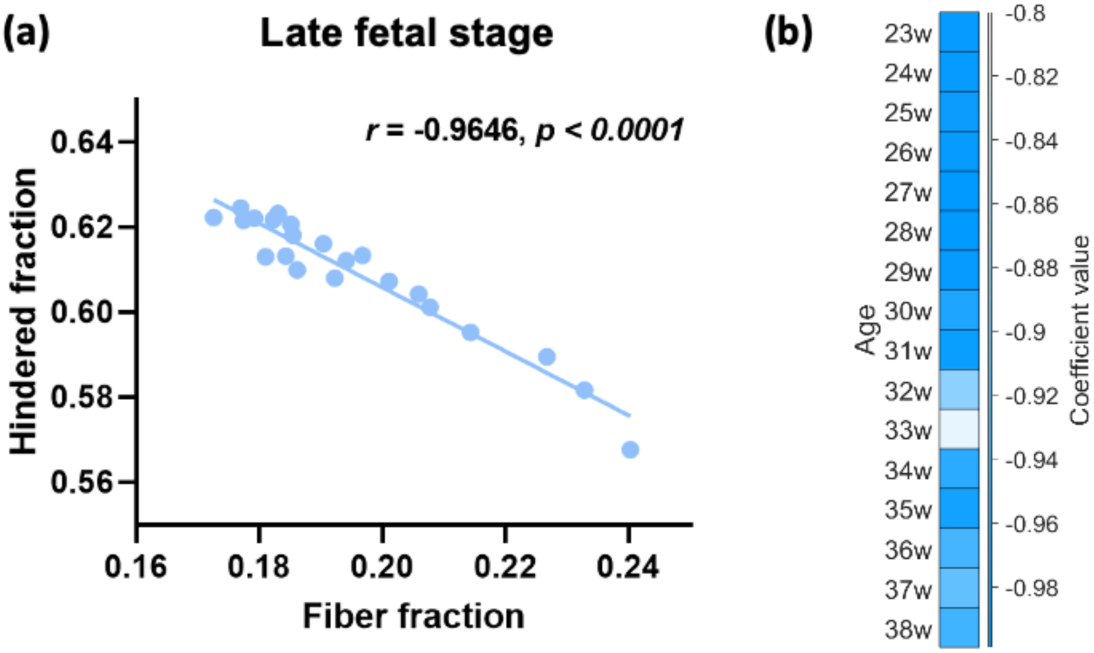
Association between DBSI fiber and hindered fractions across cortical regions. (a) An example correlation result at late fetal stages, using the same 22 labeling ROIs and gestational ages applied in the imaging-genetic analysis. (b) Consistently negative correlation coefficients across 23-38 weeks of gestation. Coefficients represent Pearson correlation betas between fiber and hindered fractions across 22 ROIs at each GA, all surviving Bonferroni correction.

**Fig. S6.**
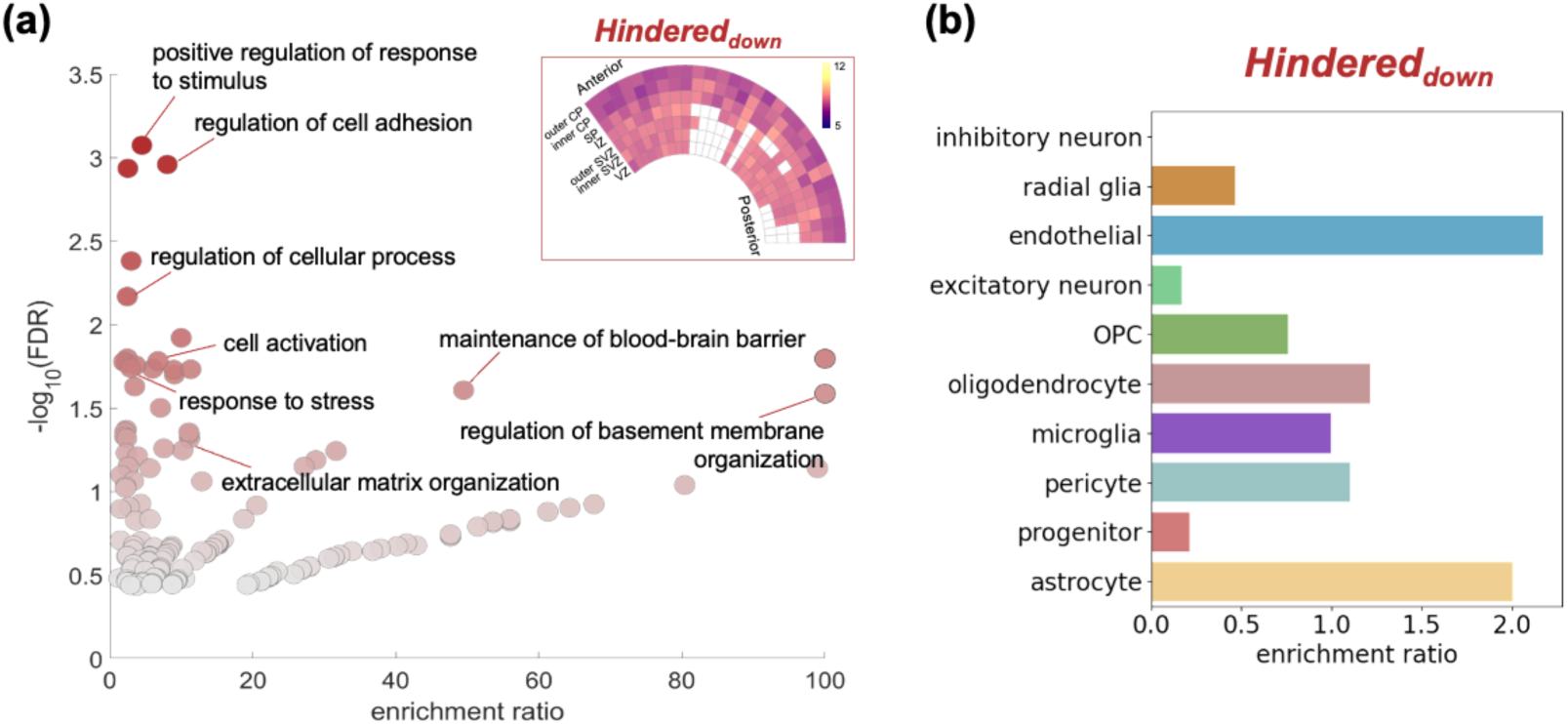
Late-gestation microstructural programs associated with cortical folding (*Hindered_down_*). **a.** Representative GO biological processes enriched in *Hindered*_down_ genes. Subplot reflects the laminar expression patterns at 21 pcw for the same gene set (rows = cortical layers; columns = ROIs ordered from anterior to posterior; box colors indicate relative (z-scored) expression level). **b.** Enrichment ratio for fetal marker genes expressed by each cell class for *Hindered*_down_ genes. Abbreviations: CP, cortical plate; SP, subplate; IZ, intermediate zone; SVZ, subventricular zone; VZ, ventricular zone; OPC, oligodendrocyte precursor cell.

**Fig. S7.**
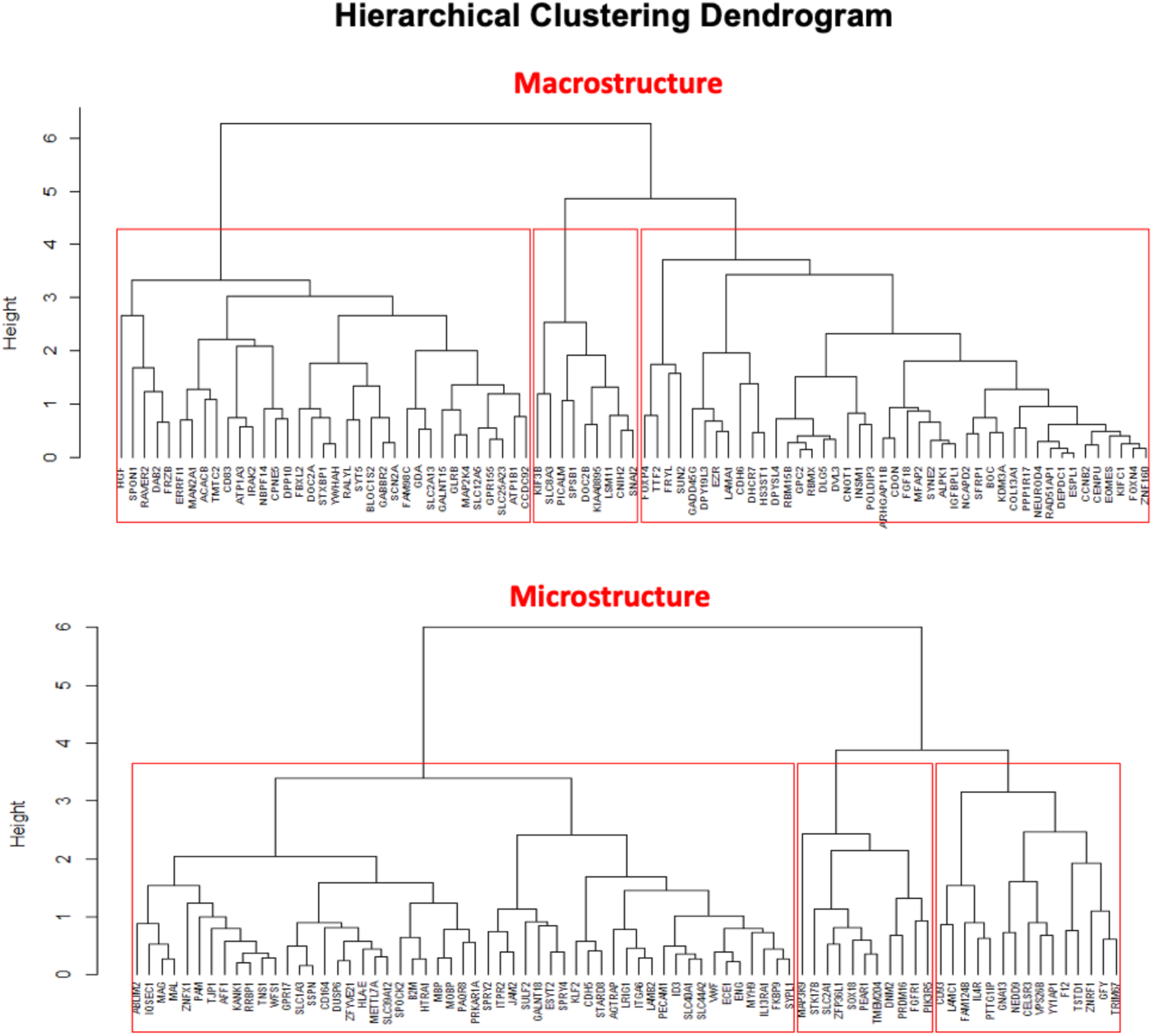
Hierarchical clustering of developmental gene expression trajectories. Unsupervised hierarchical clustering was performed to group the genes associated with macrostructural (Top) and microstructural (bottom) cortical folding phenotypes with similar nonlinear lifespan expression patterns.

## Footnotes

1 https://github.com/zjuwulab/CHN-fetal-brain-atlas

2 https://github.com/zjuwulab/Fetal-Brain-dMRI-Atlas

